# A Mechanical Model of Early Somite Segmentation

**DOI:** 10.1101/804203

**Authors:** Priyom Adhyapok, Agnieszka M Piatkowska, Michael J Norman, Sherry G Clendenon, Claudio D Stern, James A Glazier, Julio M Belmonte

## Abstract

The clock-and-wavefront model (CW) hypothesizes that the formation of somites in vertebrate embryos results from the interplay of molecular oscillations with a wave traveling along the body axis. This model however does not explain how molecular information is interpreted by cells to modulate their rearrangement into somites. Here we performed Scanning Electron Microscopy (SEM) on the pre-somitic mesoderm (PSM) of chicken embryos at stages 11-12 to describe in detail the cell shape changes occurring along the axis of the PSM. This reveals a wave of epithelialization of the dorsal PSM that precedes somite segmentation. Signs of spatially periodic apical constriction appear in this layer starting at least 3-4 somite lengths caudal to the most recently formed somite. The sizes of these clusters correspond to the typical diameter of chicken somites. We propose that a mechanical instability process leads to the separation of cells into these structures and positions the future inter-somite boundaries. We present a model in which a wave of apical constriction leads to increasing tension and periodic failure of adhesion junctions within the dorsal epithelial layer of the PSM, thus positioning somite boundaries. This model can produce spatially periodic segments whose size depends on the speed of the contraction wave (**W**) and the rate of increase of apical contractility (Λ). The Λ/W ratio determines whether this mechanism produces spatially and temporally regular or irregular segments, and whether segment sizes increase with the wave speed (scaling) as in the CW model. We discuss the limitations of a purely mechanical model of somite segmentation and the role of biomechanics along with CW during somitogenesis.

## INTRODUCTION

Somitogenesis in vertebrate development sequentially and periodically creates metameric epithelial balls (somites) along the elongating embryo body from bilateral rods of loosely connected mesenchymal cells called pre-somitic mesoderm (PSM). As cells leave the rostral/anterior (head) end of the PSM to form each somite, new cells continuously move from the tail bud to join the PSM at the caudal/posterior (tail) end of the embryo [1]. At any given rostral-caudal position, a pair of nearly equal-sized somites form simultaneously on both sides of the neural tube, between the ectoderm and the endoderm. These transient structures are the precursors of vertebrae, ribs and many skeletal muscles; many birth defects are associated with a failure in one or more steps in this long developmental process [2].

Somitogenesis is strikingly robust to perturbations (both spatial and temporal). Changes in the total number of embryonic cells or in the rate of new cell addition at the caudal end lead to compensating changes in the size and timing of somite formation so that the embryo eventually produces the same final number of somites as in normal development [3-4]. One way to achieve this conservation of final somite number is for the size of the somites to increase linearly (*scale*) with the speed of the caudal-moving position of the determination front or with the rate at which cells join the caudal end of the PSM.

Models seeking to explain somite formation include the ‘cell cycle model’, which couples the timing of segmentation to the progression of cells through the cell-cycle and a cell-intrinsic gating mechanism [5-7] and reaction-diffusion models [8-9]. Currently, the most widely accepted family of models employ a ‘clock and wavefront’ (CW) mechanism which combines caudally progressing waves of determination and differentiation with an intracellular oscillator which determines cell fate based on its phase at the moment of determination [10]. Following the identification of the first oscillating transcripts (*hairy1* and *hairy2*) in the PSM [11], many computer simulations of varying complexity have implemented different CW models. Most CW models reproduce the experimentally observed scaling of somite size with clock period, wavefront speed and rate of elongation of the PSM [12-14].

Recent experiments have shown that somite-like structures can form without either a clock or a wavefront [9,15]. The ability of somites to form without either a clock or a wavefront suggests that we should consider other mechanisms that could lead to spatially and temporally periodic sequential division of the PSM into regular segments. Moreover, recent experiments by Nelemans and colleagues showed that applied tension along the rostral-caudal axis can induce the formation of intersomitic boundaries in locations not specified by CW signaling [16], suggesting that mechanical mechanisms may be important in generating intersomitic boundaries.

In 2009, Martins and colleagues imaged the morphology of cells during chicken somitogenesis *in vivo*, showing that cells elongate, crawl and align with each other as they form a somite [17]. Importantly, their observations show that cells epithelialize gradually during somite formation, with epithelialization beginning before segments separate from each other. This finding is consistent with other reports showing that PSM cells gradually increase the density of cell-cell adhesion molecules [18] and decrease their motility [19-20] as they approach the time of the physical reorganization associated with somite formation. Moreover, several decades ago, scanning electron microscopy (SEM) observations of the PSM in a variety of species led to the suggestion that “pre-somite” like structures, named “somitomeres”, precede the condensation of cells into somites by at least 2-3 somite lengths [21-22]. However, these observations were made mostly in randomly fractured embryos along various planes and it is difficult to determine the progression of somitogenesis from them.

Here we took a new look at the PSM cells to investigate this early organization in more detail. Consistent with previous work, our observations show that dorsal PSM cells undergo an early maturation process, forming an epithelial monolayer along the ectoderm boundary, beginning long before somite formation. This pre-somitic epithelium also shows signs of a pre-segmentation process, with cohorts of cells forming arched tissue segments of roughly the length of a somite diameter.

These observations suggest that periodic tissue segmentation and somite boundary positioning could result from a mechanical instability mechanism, similar to what is seen in periodic cracking of materials subjected to stress. We developed a model of the dorsal epithelial monolayer where the observed boundaries between dorsal segments arise from the break of contact between neighboring cells due to increased apical tension between cells. We simulated this scenario with a 2D computational model of a cross-section of the epithelial monolayer and showed that spatial periodicity in segment sizes can be explained by a simple mechanical model without a clock. We also showed that this model can produce either constant-size segments or segments whose size increases (*scales*) with wave speed and inverse rate of increase of apical contractility. A roughly constant ratio of the wave speed to apical contractility build-up rate defines the boundary between these two domains. A second ratio also predicts when this mechanism produces spatially and temporally regular segments from irregular segments.

## RESULTS

### Early signs of boundary specification

To investigate the beginning of epithelialization, we performed 3D Scanning Electron Microscopy (SEM) of chick embryos fixed at various stages of somitogenesis, fractured as precisely as possible along parasagittal or transverse planes [27]. We then manually defined the contour of each individual PSM cell and calculated their aspect ratios (Figure 1A,B).

**Figure 1.**
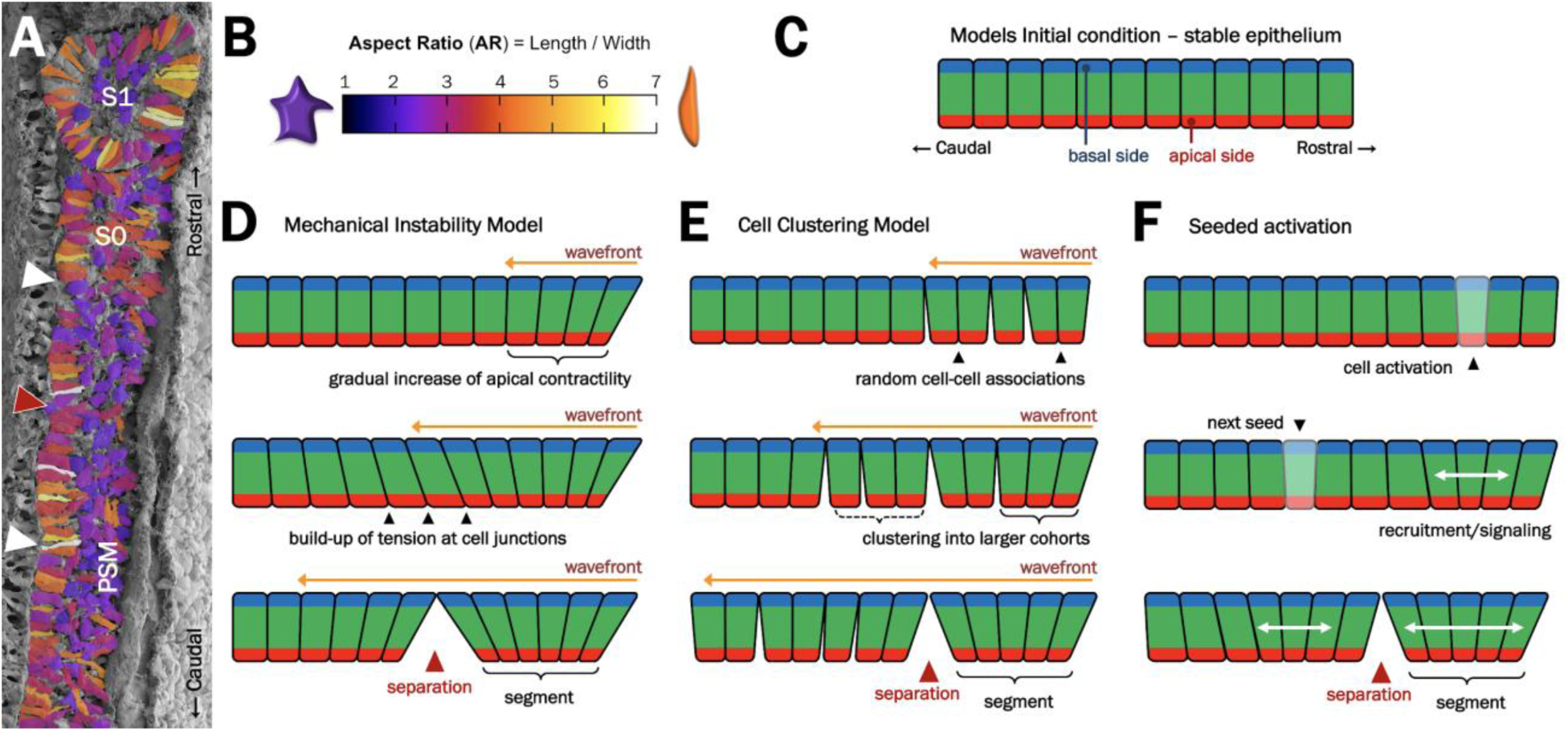
Early signs of pre-somitic epithelium segmentation and proposed models of segment formation. (**A**) SEM images of para-sagittally sectioned chicken embryos shows the epithelialization of dorsal PSM cells at least 5 somite-lengths caudal (left) to the S1 somite. Cells are colored according to their aspect ratio, as shown in (**B**). White arrowheads show the positions of segment boundaries. (**C**-**F**) Models of mechanical-instability mechanism of tissue segmentation. (**C**) Epithelialized cells, with defined apical (red) and basal (blue) sides form a rostral (right)-caudal (left) monolayer along the dorsal side of the PSM. (**D**) Mechanical instability model: a caudal-moving wavefront of myosin activation (orange arrows) initiates apical constriction of the cells in the monolayer eventually leading to the periodic segmentation of the tissue. (**E**) Cell condensation model: a caudal-moving wavefront of maturation (orange arrows) initiates random cell-cell groupings that eventually organizes the tissue into regularly sized cohorts. (**F**) Seeded activation mechanism: instead of a continuous wave, small groups of cells are activated and rearrange neighboring cells into a segment.

Our observations show that a dorsal layer of PSM cells begins to epithelialize at least as early as 4-5 somite-lengths caudal to the most recently formed somite (S1). Furthermore, cells within this dorsal epithelium form a series of cohorts at least 3-4 somite lengths caudal to the forming somite (Figure 1A). Cells near the dorsal surface within these cohorts are usually wedged shaped, with their apical (ventral facing) sides more constricted than their basal (dorsal facing) sides. The sizes of these cohorts are regular and roughly correspond to the rostral-caudal length of the future somites.

### A mechanical model of dorsal PSM pre-patterning for segmentation

The correlation between cohort size with somite rostral-caudal length and tear positioning with projected somite boundaries suggest that these structures are precursors of the future inter-somite boundary. But how do they arise? Figure 1D-F illustrates some possible mechanistic models, including a cell clustering model with a continuous wavefront (Fig. 1E) and other variations where changes in cell behaviors do not progress continuously from caudal to rostral (Fig. 1F).

Here, we take the convergence of the cells’ apical sides within the cohort as an indication that cells are undergoing apical constriction as the PSM matures. Apical constriction is the result of contractile forces generated by myosin activity at the cells’ apical side, which often bring neighboring cells together and give them a wedge shape [28-32]. It is known that when apical contractile forces exert too much load on junctional adhesion sites, tissue tears appear, as observed in *Drosophila* embryos [33-34]. We postulate that this mechanism - a mechanical instability model - may be behind the pre-segmentation pattern we observe: a caudally travelling activation wave induces apical constriction along the maturing PSM cells (Figure 1D), leading to a buildup of cell-cell apical tension (Figure 1E) that eventually subdivides the monolayer into regularly sized segments (Figure 1F). This is the model we explore in this paper.

The model has three main variables: the speed at which this caudally-travelling wave passes through the tissue (*W*), the rate at which each pair of activated cells increases its apical contractility (*Λ*), and the maximum tension a cell pair can sustain before the cells break their connection to each other (*T*_*break*_). For any given value of *T*_*break*_, we expect average segment sizes to increase proportionally with *W* and inversely with *Λ*. In the following subsections we implement two different versions of this model to explore whether and under which conditions this process can produce spatially and temporally regular segments.

### Linear elastic model of tissue segmentation

We first implemented the mechanical model of the interaction between adjacent cells in the dorsal monolayer as a series of springs connecting a chain of point objects. The model creates motion by linearly increasing the spring constant over time at rate *Λ*, with a delay between spring activation based on how far along the chain the spring is (*W*). At each time step, the acceleration due to spring forces is calculated and used to set up a system of 2*N* first order ordinary differential equations (ODEs) with time dependent coefficients, where *N* is the number of cells (points) in the system. If at any given time step the tension of a spring exceeds *T*_*break*_, the spring constant is set to zero for the remainder of the simulation, effectively breaking the spring. The time and location of each break is recorded. For more details on the model implementation, see *Computational Models*.

For short times, each cell pair tension evolves with the same patterns, first increasing roughly linearly over time after activation, until eventually turning over, decreasing for a short time and increasing again at a slower rate (Supplemental Figure 1A). During this process, the subsequent cell pair tensions follow a similar pattern, but time-delayed and reaching higher tensions before turning over. Eventually one spring reaches the threshold tension, setting the first break. We found the size (or position) of the first break, which we call the segment size (*S*), to be proportional to *W* and inversely proportional to *Λ*, as expected (Supplemental Figure 1B).

However, after the first break happens and tension is lost on one side of the monolayer, the cell centers (the point objects) begin to collapse quickly and the cell positions start to overlap (Supplemental Figure 1C). From there onwards the physical requirement that the order of the cells must be preserved is no longer guaranteed and the subsequent breaking points lose their biological meaning. Addition of spring resting lengths and ceilings on the maximum spring constant did not improve the oscillatory behavior and cell position overlaps after the first break.

### A Cellular Potts implementation of the mechanical instability model

The above results suggest that a more realistic representation of the dorsal cells that takes into account cell shapes, viscosity and thermal fluctuations, is needed for the implementation of a mechanical instability model of tissue segmentation. In the subsections that follow we implemented a stochastic Cellular Potts Model (CPM)/Glazier-Graner-Hogeweg (GGH) model version of the dorsal tissue segmentation process. (For more details see *Computational Models*) In this model, the cells are not points, but spatially extended objects similar to the schematic cells in Figure 1C-F with a width and height; apical, basal and core domains; and elastic connections between neighboring apical domains that follows the same rules as in the purely elastic model. We first characterize the model behavior with different values of a fixed spring constant (*λ*_*A*_) and simultaneous rate of increase of apical contractility on all cells (*Λ*) before exploring the gradual, caudally-moving wavefront of apical constriction activation.

### Tension profiles as a function of number of cells and constriction strength

We first test our model by creating small epithelial monolayers composed of 4 to 16 cells of fixed aspect ratio *AR* = 2 (Figure 2A). The strength of apical contractility of all cell pairs was increased simultaneously at a fixed build up rate of *Λ* = 0.05, from *λ*_*A*_ = 0 up to a maximum value of *λ*_*A*_ = 600, without allowing apical links to break. We then observe the shape of the tissue and the average cell pair tension (Eq. 6) over 20,000 simulation time units (defined in terms of Monte Carlo Steps (MCS)), after *λ*_*A*_ has reached its maximum value. As the number of cells in the monolayer increases, cell pairs in the middle of the segment are under higher tension compared to cell pairs near the periphery (Figure 2B,C). This forms the premise of our model - as the tissue becomes larger, more tension is accumulated between cell pairs, thus predisposing the tissue to break into smaller segments when a breaking tension threshold is present.

**Figure 2.**
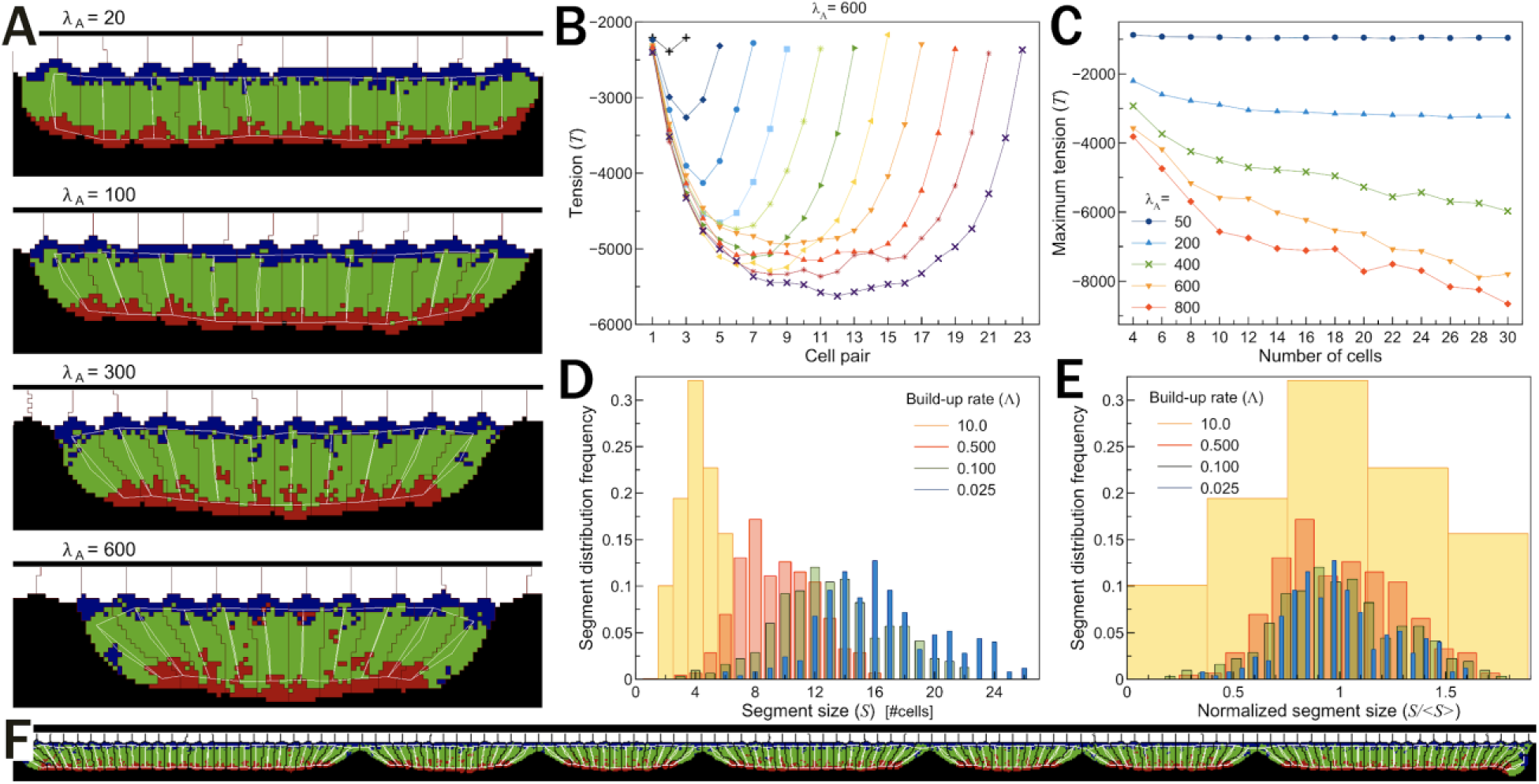
Tension and segment size distributions for fixed tissue sizes and simultaneous increase in contractility. (**A**) Snapshots of 4 simulations of 10 cells with different levels of maximal apical contractility strength. The tissue becomes more constricted with larger values of *λ*_*A*_. (White - ectodermal tissue; black - non-modeled PSM; cells domain colors as in Figure 1C-F; vertical white lines - internal distance constraints between cell domains (Eq. 3); horizontal white lines - distant constraints between domains of neighboring cells (Eq. 5)). (**B**) Plot of average apical tension (Eq. 6) between cell pairs at the end of multiple simulations with different numbers of cells (from 4 to 24). In all these simulations *λ*_*A*_ is set to 600. (**C**) Plot of maximum cell pair tension versus the number of cells in the tissue for different maximal values of *λ*_*A*_. (**D-E**) Histogram of distribution of segment sizes (**D**) and normalized segment sizes (**E**) for different build-up rates of apical contractility. (**F**) Snapshot of a simulation with *Λ* = 0.05 showing a wide distribution of segment sizes.

### Simultaneous activation of apical contractility leads to irregularly sized segments

Next, we combine the simultaneous activation with the breaking of the apical links by imposing a tension threshold *T*_*break*_ = −7500 after which the link between the apical domains of neighboring cells is broken. As in the previous section, in these simulations the strength of apical contractility is increased linearly from *λ*_*A*_ = 20 up to a maximum value of *λ*_*A*_ = 600 at different rates of apical contractility build-up (*Λ* = *dλ*/*dt*). Since now any cell contact is initially equally susceptible to break, we obtain a broad distribution of segment sizes (Figure 2D-F), with the average segment size increasing as the build-up rate decreases (Figure 2D). The shape of the distribution, however, remains the same for all values of *Λ* (Figure 2E), except when the rates are bigger than 10, where we found that most cell pairs break. The data in Figure 2D-F are from simulations with periodic boundary conditions, but the results are qualitatively the same for simulations with a large number of cells (*N* > 400) and fixed boundaries (Supplemental Figure 2). We conclude that a simultaneous activation of constriction activity is insufficient to produce a regular pattern of pre-segments similar to those seen in our SEM observations of embryos (Figure 1A).

### Gradual and sequential activation of apical contractility leads to tissue segmentation

Next, we investigated whether sequential activation of apical constriction can slice a tissue to generate regularly sized segments. From now on, all simulations have a large number of cells and a caudally moving front of activation that sequentially initiates a gradual and linear increase of the strength of apical contractility of cell pairs. The wave speed *W* and build-up rate of apical contractility *Λ* will be varied systematically around their base values (see Table1). In addition, apical cell pairs will have a maximum tension load *T*_*break*_, after which their links break and a tear may appear between the apical region of those cells.

Figure 3A shows a time series of a zoomed-in section of a simulation with 115 cells where 4 tissue segments form. With the exception of the first and last two segments (due to boundary conditions), all segments are of similar size (< *S* >= 11.36 +/− 1.45) and segmentation occurs at similar time intervals (< *τ* >= 3442.37 +/− 587.87) for our standard set of parameter values (see Table1). We also looked at the evolution of the tension profile for 2 sequential segments. As the wavefront of activation passes, the tension between cell pairs gradually increases, with the rostral-most pair with the higher tension (Figure 3B and green lines in Figure 3C). After the formation of the rostral-most boundary/tear the pattern inverts, with the rostral-most cell pairs now more relaxed and the caudal-most pairs now under higher tension (compare green and blue lines in Figure 3C). Formation of the caudal-most boundary relaxes the tension of these pairs and the segment tension profile has now a symmetrically convex shape with an average lower tension compared with the intermediate steps (black lines in Figure 3C). The same process repeats itself in the formation of the subsequent segments.

**Figure 3.**
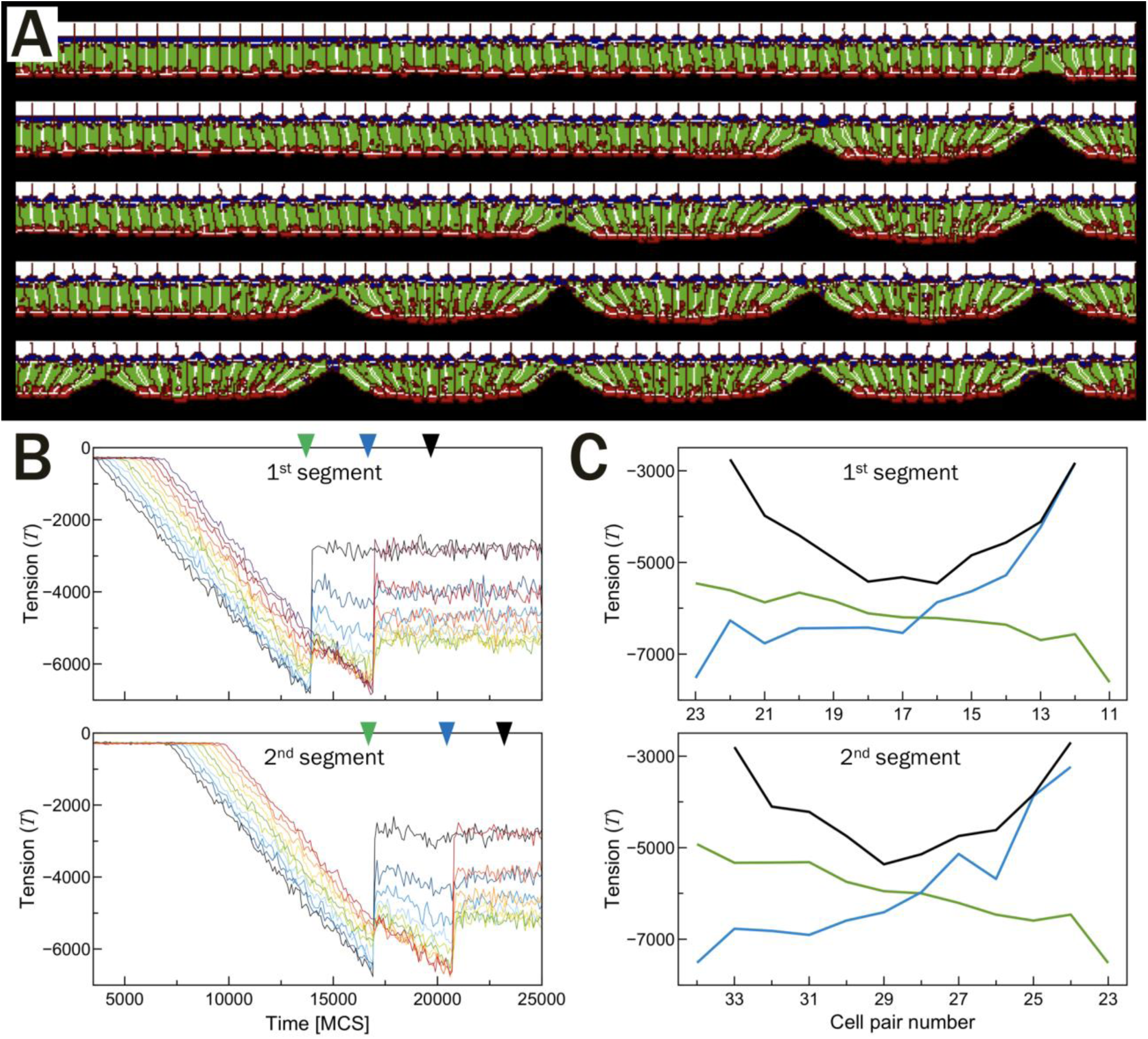
Tissue segmentation from a caudally propagating wave initiating apical constriction. (**A**) Time-series of a simulation showing the sequential segmentation of a tissue due to a linear increase of apical cytoskeletal activity as cells are progressively activated from rostral (right) to caudal (left). Colors as in Figure 3A and parameters are from Table 1. Snapshots taken at the approximate moment the separation occurs (17000, 21000, 24000, 28000 and 31000 MCS). (**B**) Time evolution of cell pair tensions in the regions of 2 consecutive segments. Lines are plotted beginning with the rostral-most pair of cells for each segment. (**C**) Tension profiles for the 2 tissue segments shown in (**B**). Each line corresponds to the same-color arrowheads in (**B**) and indicate three different events - formation of the rostral segment boundary (green), formation of caudal segment boundary (blue), and some time after the caudal event occurs (black).

### Segment size scaling with wave speed and rate of apical constriction

The clock and wavefront model was proposed to explain how somite size can adjust to variations of embryo size [10]: all else being equal, a faster clock produces smaller somites, while a faster wavefront generates larger somites. We now investigate if our mechanical model of segmentation has the same scaling features: does a faster wave of activation (*W*) lead to larger segments? How does average segment size change with different build-up rates of apical contractility (*Λ*)? In the results that follow we systematically varied both parameters and all data points were averaged over 5 simulation replicas.

**Table 1.** Reference values of the 4 key parameters of our model.

| Parameter | Symbol | Base value (units) |
| --- | --- | --- |
| Wave speed | $W$ | 0.003 (cells/MCS) |
| Apical contractility build-up rate | $\Lambda$ | 0.05 (MCS<sup>-1</sup>) |
| Apical tension breaking threshold | $T_{break}$ | -7500 (dimensionless) |
| Cell aspect ratio (height/width) | $AR$ | 2 (dimensionless) |

We first fixed the build-up rate of apical contractility *Λ* and varied the wave speed *W*. We observe two regimes of the average segment size < *S* > with respect to the wave speed. For wave speeds below a critical value (< *W*^*\**^) average segment size < *S*^*\**^ > was constant, but increased as a power law with exponents close to ¼ for faster wave speeds (> *W*^*\**^) (Figure 4A). The critical values of the wave speed *W*^*\**^ and average segment size < *S*^*\**^ > for the change from constant segment size to the scaling regime depends on the value of the build-up rate *Λ* (Figure 4B). These results suggest that the discrete spatial compartmentalization of the tissue in cell units poses a lower limit on segment size as a function of *Λ* (Figure 4B). In fact, our simulations in absence of a wave already indicate that the maximum tension a cell pair can reach depends on the number of cells within a forming segment (Figure 2B), so for slow waves we expect the tears to always occur at regular size intervals.

**Figure 4.**
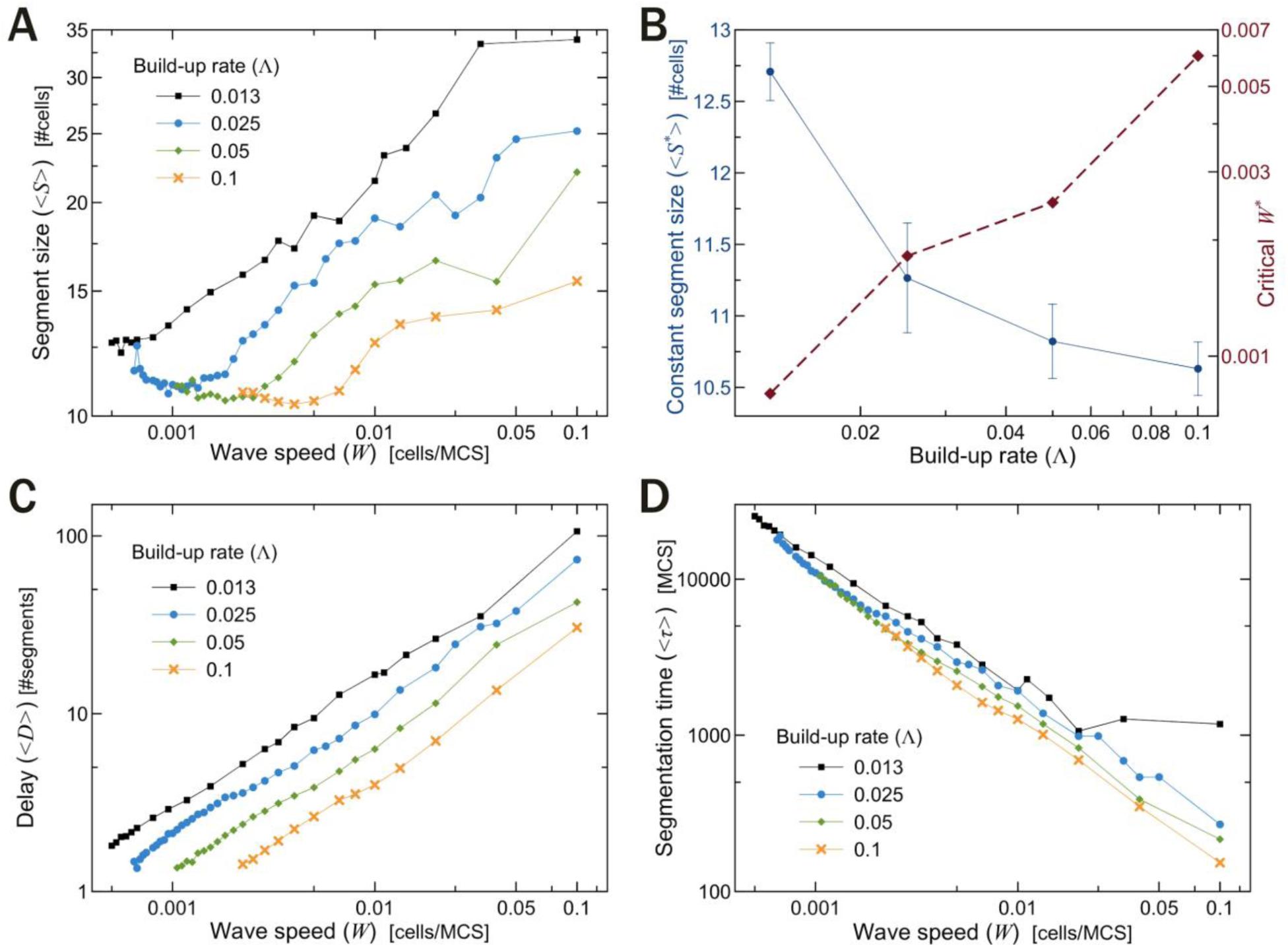
Segmentation as a function of wave speed. (**A**) Average segment size < *S* >as a function of wave speed *W*. For slower wave speeds segments are of roughly constant size, but increase as a power law for faster wave speeds. (**B**) Constant segment sizes < *S*^*\**^ > (blue solid line) and critical wave speeds *W*^*\**^ (red dashed line) as functions of *Λ*. (**C**) Distance between activation and boundary formation < *D* > increases linearly with *W*. (**D**) Average segmentation time < *τ* > decreases as a power law with *W*^−0.80±0.008^. (**A**,**C**,**D**) Each line shows a similar behavior for different base values of the build-up rate of apical contractility (*Λ*).

The average distance between apical constriction activation and boundary formation < *D* > showed a less surprising behavior with an approximate linear scaling with wave speed (Figure 4C): the faster the wave, the larger the distance between the wavefront and the boundaries. Changes in the base value of *Λ* only shifted the curves, with lower build-up rates increasing the distance between activation and boundary formation. The average time interval between the formation of successive boundaries < *τ* > depends only on *W*, and not on *Λ*, with faster wave speeds decreasing the segmentation time as a power law with exponent −0.80 ± 0.008 (Figure 4D).

Next we fixed the wave speed *W*and varied the build-up rate of apical contractility *Λ*. Average segment sizes < *S* > decrease logarithmically with higher build-up rates, but became constant after a critical value of *Λ*^*\**^ (Supplemental Figure 3A,B). As before, segment sizes outside the scaling regime (< *S*^*\**^ >) depends on the value of the wave speed *W* used in the simulations (Supplemental Figure 3B). Again, these results can be understood in light of the way cell pair tension increases with segment size (Figure 2B). For slow build-up rates (< *Λ*^*\**^), the sole factor determining segment size is the wave speed, with faster waves adding more cells to the forming segment before the boundary formation (Figure 4A). For higher build-up rates (> *Λ*^*\**^), however, the rate of cell addition is not fast enough to overcome the increase in tension profile, which itself is a function of tissue size (Figure 2B).

The average distance between cell activation and boundary formation < *D* > decreased linearly with the build-up rate, as expected: the faster the contractility build-up rate, the less time it takes for segments to form once all cells are actively constricting (Supplemental Figure 3C). The average segmentation time < *τ* > was similarly dependent on the build-up rate as on segment size < *S* >, which was expected, given that < *S* >= *W* < *τ* > (Supplemental Figure 3D).

### Ratio of build-up rate to wave speed sets the transition of scaling regimes

The critical values of wave-speed *W*^*\**^and build-up rate of apical contractility *Λ*^*\**^ for the transition between constant and segment size scaling regimes shown in Figures 4B and Supplemental Figure 3B are related. Rescaling of the horizontal axis in Figures 4A and Supplemental Figure 3A for each curve by their corresponding values of *Λ* and *W*, respectively, shows that in both cases, the transition occurs around the same ratio of *Λ*/*W* = 22 *MCS*^2^/*cell* (Figure 5A,B). This allows us to define a boundary transition in parameter space that separates regions where segment sizes change with variations in either *Λ* or *W* from regions where the segment sizes remain relatively constant with changes in one of these parameters (Figure 5C, green and blue regions). Note that we use the word *constant* in contrast to the size scaling of the segments with respect to either *Λ* or *W*, there is still a small and gradual change in segment sizes within the green region in Figure 5C (see Supplemental Figure 4).

**Figure 5.**
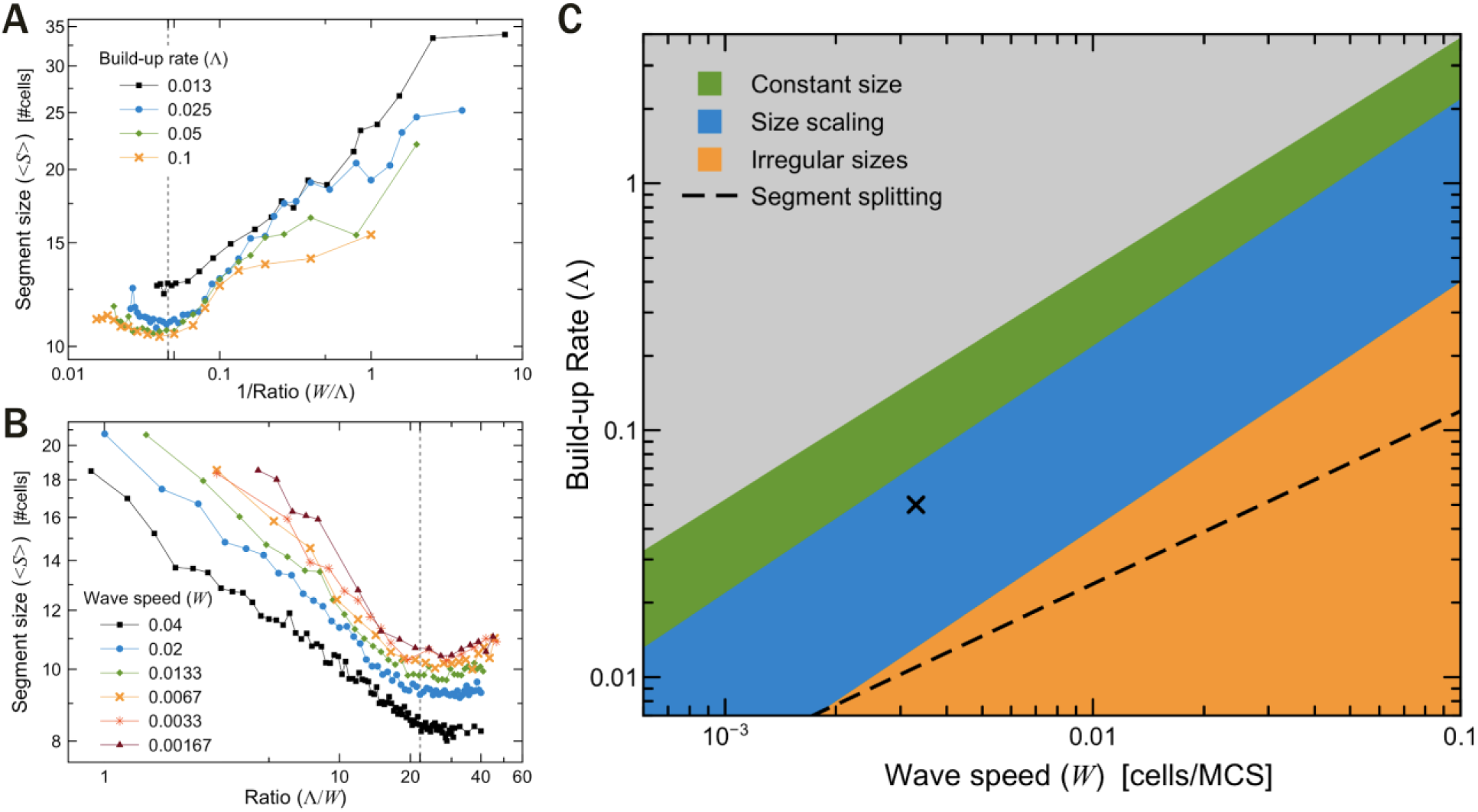
*Λ*/*W* sets boundaries between segmentation regimes. (**A**,**B**) Rescaled version of Figures 4A and Supplemental Figure 3A, showing average segment size < *S* > as a function of (**A**) wave speed to build-up rate, and (**B**) its inverse. Vertical dashed line at *Λ*/*W* = 22 indicate transition threshold between constant size and scaling segment sizes. (**C**) Parameter space diagram showing regions where the combination of the parameters *Λ* and *W* leads to constant segment sizes (green), scaling segment sizes (blue), or irregularly sized segments. Black dashed line shows transitions from stable to splitting segments. Black **X** point indicates the reference simulation parameters (see Table 1). Grey region indicates parameters combinations that lead to simulation artifacts.

The parameter space in Figure 5C also contains a grey region where the combinations of *Λ* and *W* leads to simulation artifacts (among other things, simulated cells decrease their height by around 10% and neighbouring apical compartments separate from each other without breaking their apical links). All data from this region were discarded in our analysis. The orange region and dashed boundary line will be defined momentarily.

### Segment sizes become irregular for low ratios of build-up rate to wave speed

While average segment size increases with higher wave speeds (*W*) or lower build-up rates of apical contractility (*Λ*), segment sizes distribution becomes irregular for lower ratios of *Λ*/*W* (Figure 6A-D). This can be visualized in plots of segment size variation, calculated as the ratio of standard deviation (*σ*) to the mean (< *S* >) (Figure 6E,F). When the wave speed is much faster than the build-up rate (*W* ≫ *Λ*), a large group of cells starts to constrict at about the same time. In this situation, the memory of the last boundary position is erased, and new tissue tears are more likely to appear anywhere in the tissue.

**Figure 6.**
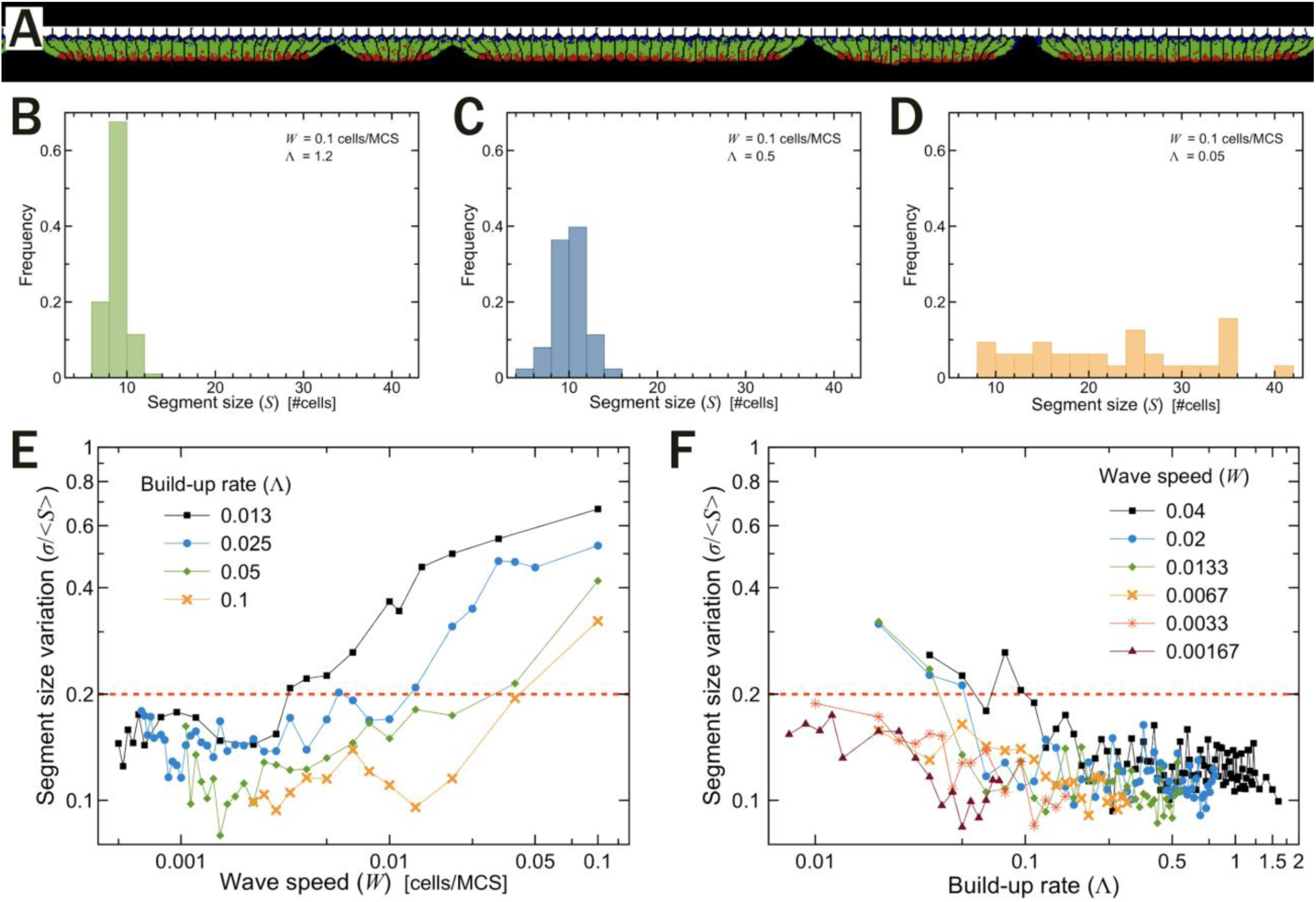
Segment size variation. (**A**) Typical simulation output showing wide range of segment sizes from the irregular region (orange) of the parameter space in Figure 5C with *W*=0.03, *Λ* = 0.013. (**B**-**D**) Histogram of segment size distributions for combinations of *Λ* and *W* in the constant size region (**B**), scaling region (**C**), and irregular region (**D**), of the parameter space over 5 trials. (**E**) Segment size variation (std/mean) as a function of build-up rate of apical cytoskeletal activity *Λ*. (**F**) Segment size variation as a function of wave speed *W*. (**E**,**F**) Dashed red lines at *σ*/< *S* >= 0.2 indicate the threshold used to determine the region of irregular segment sizes in Figure 5C.

We choose to classify segments as irregular when *σ*/< *S* > is higher than 0.2. This corresponds in our parameter space diagram, to a transition from uniform to non-uniform segments at ratios of *Λ*/*W* ≤ 4. A similar dependence on *Λ*/*W* is observed with the segmentation time variation (*σ*/< *τ* >), which measures how uniformly distributed is the time between consecutive boundary formations (see Supplemental Figure 5). A threshold of *σ*/< *τ* >=0.33 can be similarly used to distinguish between regular and irregular segments. This high variation in segmentation time in the irregular region is expected, as the time taken to form subsequent tears is an emergent output from our mechanical model, rather than a direct input related to the oscillation period of the clock as in the CW model.

### Build-up rate and wave speed limit larger segment sizes

The above results show that initial segment sizes increase with faster wave speeds (Figure 4A) and slower build-up rates of apical contractility (Supplemental Figure 3A). However, there seems to be a limit on the larger segment sizes. After their initial formation, larger segments are prone to split as the cells within them continue to increase their apical cytoskeletal activity strength (*λ*_*A*_) and tension keeps building up until a new splitting event occurs near the middle of the formed segment (Figure 7A,B).

**Figure 7.**
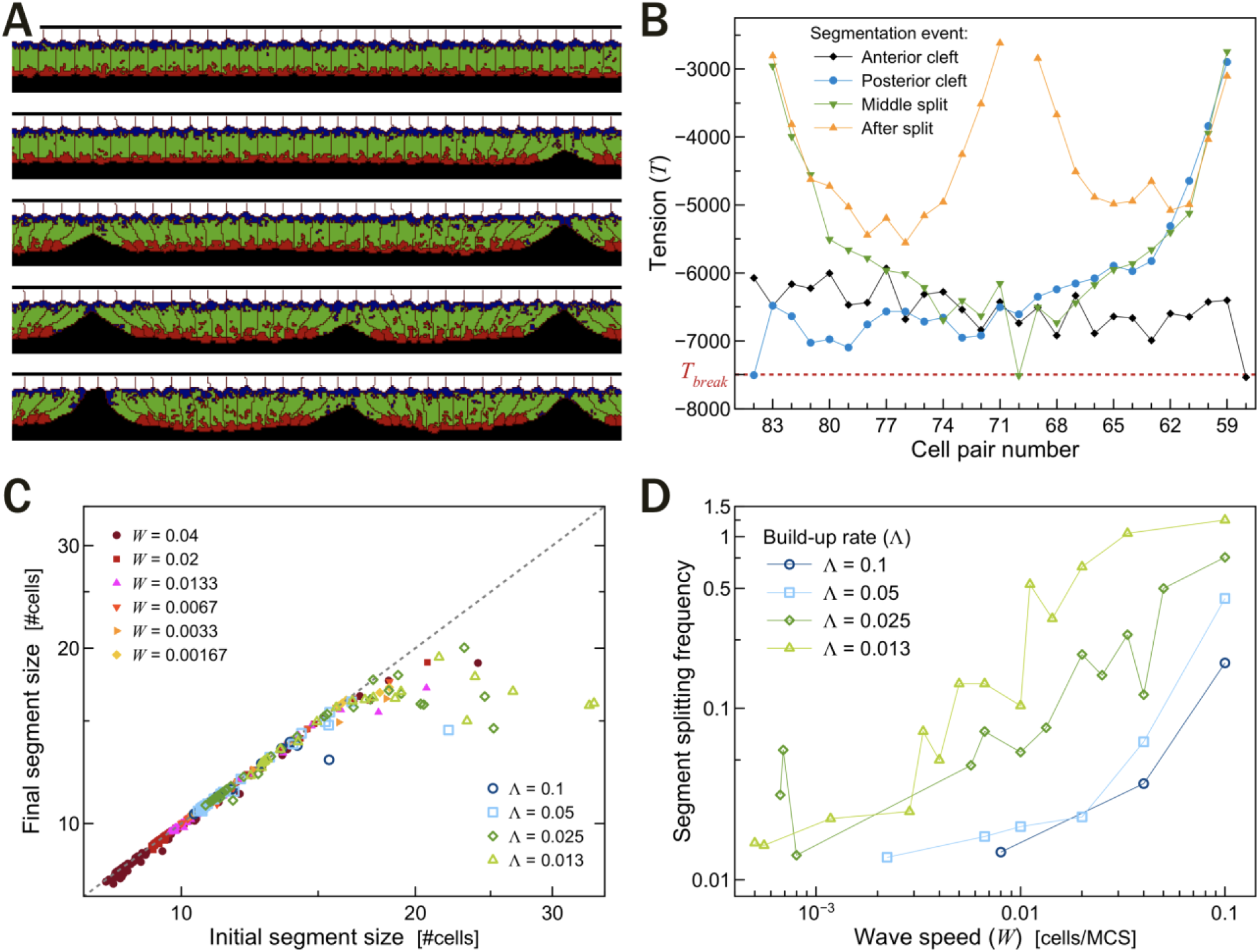
Splitting of Large Segments. (**A**) Simulation snapshots showing the splitting of a large segment (times, from top to bottom: 47000, 49000, 52000, 54000 and 56500 MCS). (**B**) Evolution of the tension profile for the splitting segment shown in (**A**). (**C**) Plot of average final versus initial segment sizes. Data points at the diagonal gray slashed lines indicate no splitting events. (**D**) Splitting frequency as a function of wave speed *W*.

While the occurrence of a splitting event is proportional to the initial segment size (Supplemental Figure 4), the wave speed *W* has a more direct effect on the frequency of splitting, with higher build-up rates only delaying the appearance of splits (Figure 7D). Together these results suggest that our mechanical model of segmentation sets a higher limit on segment sizes depending on the speed of the wave with respect to the build-up rate of apical cytoskeletal activity (Figure 7C,D). Our results, however, did not point out to a specific ratio of *Λ*/*W* as predictive of a transition to a higher frequency (>0.1) of splitting events, but rather a nonlinear function *Λ* = 0.6 *\* W*^0.7^ for the boundary between stable segments and splitting segments (dashed line in Figure 5C).

### Segment size as a function of maximum tension load

So far, we had used a fixed apical to apical tension breaking threshold of *T*_*break*_ = −7500. We now test the effects of varying the threshold tension at which the link between the neighboring cells’ apical domains break. At the extreme limit of |*T*_*break*_| near zero, the average segment contains only one cell. Conversely, at high enough values (|*T*_*break*_| > 9500) the tissue never segments and remains a single monolayer. As the magnitude of the breaking threshold increases between these ranges, the average segment sizes also increase, as expected (Figure 8A). Note however, that the splitting frequency is higher for low values of |*T*_*break*_|, and zero for high values, with segments breaking in more than once for threshold values around *T*_*break*_ = −2000 (Figure 8B).

**Figure 8.**
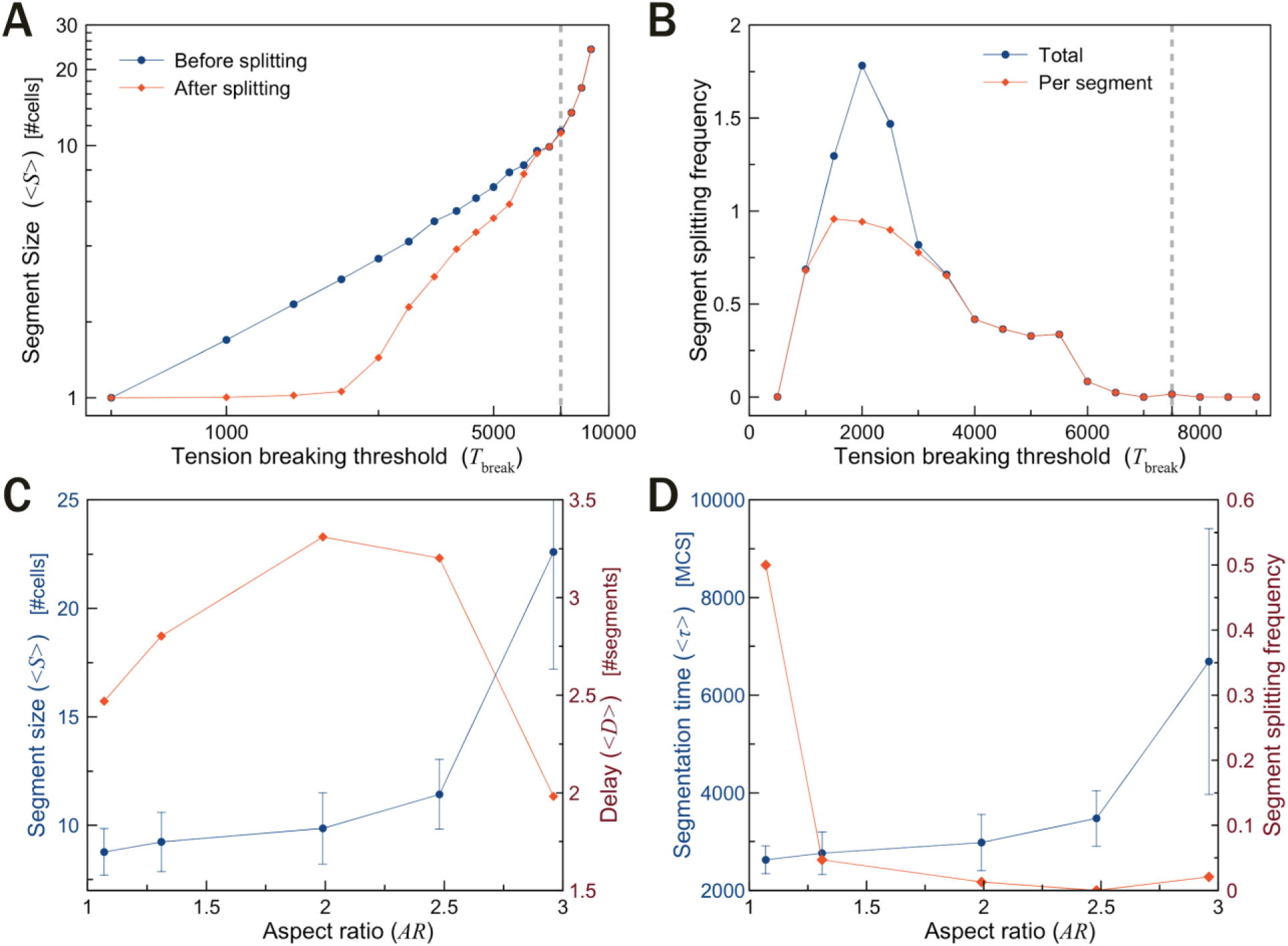
Influence of junctional tension limit and cell shape on segmentation. Average segmentation size < *S* > as a function of the absolute value of tension breaking point |*T*_*break*_ |. Blue lines are initial segment sizes, red lines are segments sizes after splitting. (**B**) Average number of splitting events per segment as a function of |*T*_*break*_ |. (**C**) Average segmentation size < *S* > (blue) and the distance between activation and boundary formation < *D* >(red) as functions of cell aspect ratio *AR*. (**D**) Average segmentation time < *τ* > (blue) and frequency of segmentation splitting (red) as functions of cell aspect ratio *AR*.

### Segmentation as a function of cell shape (aspect ratio)

In the pre-somitic tissue, cells elongate dynamically prior and during the process of boundary formation. We had kept the length of the cell fixed for our previous simulations and we now evaluate how segment size depends on the aspect ratio of the cell (*AR* = *height*/*width*). Figure 8C shows the average segment size < *S* > and distance < *D* > as a function of cell aspect ratio. As expected, it is easier to pack more elongated cells into a segment. The distance < *D* >, however, is a concave curve with respect to *AR* with a maximum around *AR* = 2. This can be understood as follows: the wider the cells, (*AR* < 2) the harder it is to pack them into a segment made of wedge-shaped cells, so apical tension builds up quickly and tissue separation occurs sooner, thus reducing both the segmentation time (blue line in Figure 8D) and the distance between activation and boundary formation compared to the standard case (Figure 8C). Conversely, the more elongated cells (*AR* > 2), the greater the number of cells that can be packed into a segment and the longer it takes for consecutive boundaries to form. However, since the segments themselves are larger, < *D* >, which is calculated in terms of segment sizes, also decreases. Figure 8D also shows that the segment splitting frequency (red line) is significantly higher for low aspect ratios, which can be understood as a result from the packing constraints that expedites the accumulation of tension between apical cell pairs and make them more prone to splitting.

## DISCUSSION

In this paper, we describe a mechanical model of tissue segmentation where a continuous wave of apical constriction activation coupled with maximum apical tension load sets the position of future somite boundaries. The model is based on experimental observations of extensive cell rearrangements during somite formation [17], observations with somitomeres [21-22] and our own SEM images (Figure 1A) suggesting that the segmentation process begins dorsally in the PSM at least 4 somite lengths prior to the formation of the last somite. We interpreted the spatial segmentation of the dorsal epithelium into similarly sized cohorts of cells as the result of an increased longitudinal (rostral-caudal) tension between the cells.

A similar process is known to cause the appearance of periodic cracks on non-biological thin films subjected to tensile stress [35-36]. Analysis of these films show that the spacing between these cracks depends on the stress, thickness and toughness of the film. In these models, however, the stress is usually assumed to be uniform, non-time varying, and applied externally to the material. One exception is the work by Yuse and Sano, where a time-varying inhomogeneous thermal stress was applied to a plate moving between two reservoirs [37-38]. The velocity of the plate and the temperature difference between the two reservoirs act as control parameters and define a phase space where periodic cracks could transition into other morphologies. Our model on pre-somitic tissue segmentation differs from these analogies in two ways, firstly, the material is ductile, and secondly and mostly importantly, the material is not homogeneous, but subdivided into discrete units (the cells).

In biology, mechanical forces generated by fibroblasts have been shown to assist in the remodeling of the external collagen matrix to form self-organized tissue structures [39]. Tensile stresses generated by stretching fibroblasts suspended in collagen and restrained by a glass mesh have also been known to result in aggregations of cells. The self-organization of cells into clusters in the absence of chemical signals led Harris and colleagues to suggest that mechanical instabilities could act as sources of positional information instead of diffusible factors like morphogens [40]. More recently the possibility of mechanical instabilities resulting in vertebrate segmentation has also been explored by Truskinovsky and colleagues [41]. Assuming relaxation of junctional adhesion sites as the PSM elongates, the model identifies that the number of somites generated by their pre-patterning mechanism is robust to the final segmentation process. Their static model, however, neglects the sequential development of the boundaries and does not take into account any active processes within the tissue.

Given these differences we developed our own model of epithelial segmentation where we identified four key parameters: the speed of the wave (*W*), the rate of increase of apical contractility (*Λ*), the maximum apical cell-cell junction load (*T*_*break*_), and the aspect ratio of the cells (*AR*) (Table 1). While the parameters have been chosen to imitate segment size in chicken, we comment that we need better tuning of the aspect ratio of the cells in the dorsal region, where we found that cells can reach aspect ratios as high as 7. To compare results at these higher aspect ratios we would require re-parametrizing our cell volume to allow for sufficient apical surface so that the width is much greater than a chosen *L*_*AT*_. While these changes are easily programmable, they come at a greater computational cost.

Other possible explanatory models include a cell clustering model (Figure 1E). Here we assume that the dorsal cells are initially confluent, but not tightly connected at their apical sides. As a wavefront of maturation passes, cells start to connect with their neighbors and condense initially into small groups, later clustering into larger, more tightly connected cohorts that eventually pattern the dorsal tissue into distinct tissue segments. Alternatively, instead of a continuous wavefront of activation/maturation, both processes (apical constriction and cell clustering) can be initiated at the center of the future segment by a small group of cells that eventually recruit and/or activate their neighbors (Figure 1F). This mechanism of initiation still assumes some kind of caudally moving wavefront, but also requires some positioning mechanism that may be provided as an earlier clock read-out. All those models, including the mechanical instability model explored here, provide an initial segmentation pattern that would later guide the formation of the full somite as the epithelialization process spreads to the ventral, medial and lateral sides of each forming epithelial sphere. These processes assume some kind of wavefront but are in principle independent of a clock, which is necessary for the molecular patterning of the somites into caudal and rostral identities [15].

We found that our purely mechanical model (Figure 1D) is able to produce spatially and temporally regular segment sizes. Similar to the CW models, segment sizes scale (increase) with higher wave speeds as seen in many species (Figure 4A). We found, however, that this scaling is not linear with wave speeds, as would be expected from a simple deterministic version of the CW model. There are two limits to this scaling behavior, which can be characterized by *Λ*/*W*ratios alone (Figure 5C). An upper limit in segment sizes is reached for low ratios (*Λ*/*W* < 4), where initially formed segments are prone to splits due to mechanical instabilities; and a lower limit in the sizes is reached for high ratios (*Λ*/*W* > 22), where segment sizes do not scale with wave speeds. These limits in somite size are absent in CW models and offer an explanation for the observed splitting of large somites formed in some perturbation experiments [42-43]. Segmentation time in our model is not imposed but rather calculated as the difference in time for the appearance of the rostral and caudal boundaries. We find that it decreases with wave speeds and is insensitive to build-up rates (Figures 4D, 5D).

Our model predicts that disruptions in the protein levels of molecules (eg myosin, actin) that control the apical contractility of the PSM cells will affect the distributions of segment sizes. Upregulating these molecules will have a similar effect as increasing *λ*_*A*_ (Fig 2C), leading to smaller segments. However, since the segment sizes in our model is also dependent on the sequential rostral-caudal activation of these molecules, we expect cases in which for a fixed tension load, segment sizes could become independent of the protein activity (similar to constant segment sizes in Fig 5C). Any disruptions on the strength of the adhesion junctions between the PSM cells (equivalent to lowering *T*_*break*_ in our model) will also lead to smaller segments. We also comment that experiments with a continuous growing tissue as in [41] will differ from a fixed tissue model, as in the former case we will expect a reduction in the strength of apical links *λ*_*A*_, leading to larger segment sizes.

Our model seems to have higher variations in the distribution of the segment sizes and formation time than existing mathematical implementations of the clock-and-wavefront model [14]. In a real 3D scenario however, epithelialization along the dorsal sides does not happen along a line as in our model, but within a plane that also extends along the medial-lateral direction. Addition of these neighboring cells into our model would likely reduce variation in segment size and segmentation time.

While our mechanical model drastically differs from the CW family of models with respect to the absence of an intracellular molecular clock within prospective somite cells, it still assumes the presence of some sort of caudally travelling wave, which does not need to be the same as the one postulated by the CW. We are agnostic about the nature of this wave, which can be either a cell-autonomous maturation process (starting from their addition to the tail-bud or a similar event), a tissue level processes, such as a read-out of FGF and RA levels, or a combination of both [44]. As is often the case in biology, we speculate that mechanics and molecular based oscillatory signaling may work together to yield a robust morphological outcome.

Our current model, however, is too simple to explore such scenarios as it is restricted to a 1D view of the dorsal monolayer and is aimed to reproduce segmentation events prior to the formation of a full somite. Further developments of this basic model will include the self-organization of the PSM cells ventral to the dorsal monolayer and the expansion of the model to 3D. It is known that in chicken [17] different regions of the PSM epithelialize at different times, and it would be interesting to explore the implications of these observations in an augmented version of the mechanical model presented here.

## COMPUTATIONAL MODELS

### Linear elastic model

We implement the interaction of adjacent cells as a series of springs of time-varying strength *λ*_*i*_(*t*) and resting length *x*_0_, connecting a chain of *N* point objects. While the number of cells is finite, we set it large enough (*N* = 50) such that in the time scale we consider, there are no significant edge effects and the chain is effectively infinite. Our model creates motion by linearly increasing the spring constant over time at a fixed rate (*Λ*) from a minimum value *k*_0_ to a maximum value *k*_*max*_, with a delay (or wave speed *W*) based on how far along the chain the spring is. At each time step, the acceleration due to spring forces is calculated and used to set up a system of 2*N* first order ODEs with time dependent coefficients. This time dependence means there is no practical way to solve this system analytically and we use MATLAB Ode45 to solve numerically how the cells interact over time. This is the simplest way to model this behavior since there are no random elements, and therefore this model cannot account for possible effects of viscosity or thermal fluctuations. However it means that the results we do get are precise and replicable.

Along with positions and velocities, tension in each spring is tracked over time. At each step, the tension of each spring is checked. If the spring exceeds a predetermined threshold tension (*T*_*break*_), the spring constant is set to zero for the remainder of the calculation, effectively breaking the spring. The time and location of each break is recorded.

At any given time, the acceleration of a point *x*_*i*_ connected by two springs is given by:

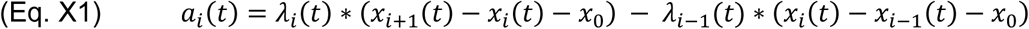

Here *t* is time, *i* is the cell position number, and *a*_1_ = *a*_*N*_ = 0, fixing the endpoints. Fixing the end could theoretically cause edge effects but the chain is long enough so that that first break always occurs before any wave behavior could reflect back into the system.

The spring constant and tensions at each cell neighboring pair are:

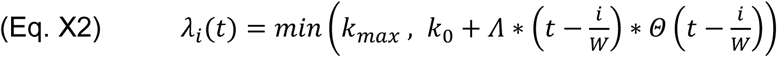

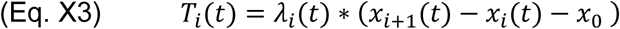

Where *Θ* is the Heaviside function.

### CPM/GGH model

We implement our full model as a simulation using the Cellular Potts (CPM), or Glazier-Graner-Hogeweg (GGH) model [23] written using the open-source CompuCell3D simulation environment [24]. The CPM/GGH framework represents each cell, or cell region as a collection of voxels that forms a domain within a fixed grid (here, square). An effective energy defines cell/domain properties such as size, mobility, adhesion preferences and distance constraints with other domains:

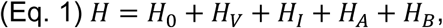

where we defined each term below.

The initial configuration of all cells/domains evolve in time by a series of random neighboring pixel copy attempts whose acceptance is governed by a Metropolis algorithm, with a number of pixel-flips equal to the grid size defining the time unit of the simulation, a Monte Carlo Step (MCS).

Similar to what has been done in previously models [15,25], each cell is composed of 3 domains representing the apical, basal and core regions of an epithelial cell. The size of the domains is maintained by a volume constraint in the Hamiltonian:

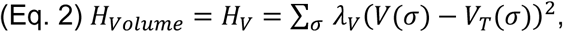

where the sum is over all domains (*σ*), *V*(*σ*) is the current cell volume, *V*_*T*_(*σ*) is the cell target volume, and *λ*_*V*_ is a lagrangian multiplier setting the strength of the constraint.

We defined the cell aspect ratio *AR* to be the ratio of apical/basal length to cell width. This ratio is maintained through spring-like distance constraints between the centers of mass of the three domains belonging to each cell:

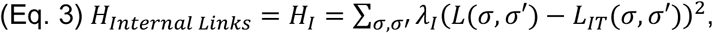

where the sum is taken over the three pairs of domains within each cell, *L*(*σ, σ*′) is the current distance between the center of mass of the two domains, *L*_*IT*_(*σ, σ*′) is the corresponding target distance, and *λ*_*I*_ is the strength of the constraint. To prevent cells from bending we set the target distance between apical and basal domains equal to the sum of the target distance between the core domain to the apical and basal domains.

Adhesion between cells is modeled with the standard Potts model internal energy term:

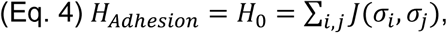

where the sum is taken over all fourth-order neighboring pairs of grid coordinates *i* and *j*; *σ*_*i*_ and *σ*_*j*_ are the cell domains at grid coordinates *i* and *j*, respectively; and *J* is the contact energy per unit area between those domains.

Apical constriction is a cell autonomous process that may lead to tissue-level events, such as invagination, through the coupling of the internal contractile activity of actin-myosin cytoskeleton of neighboring cells with their adhesion junctions. Since we are interested only on the tissue level effects of this process, we model junctional adhesion and apical constriction similar to (Eq.

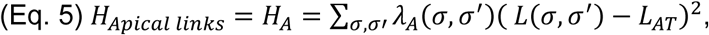

where the sum is taken over all pairs of neighboring apical domains, *L*(*σ, σ*^′^) is the current distance between their center of mass, *L*_*AT*_ is the target distance between them, and *λ*_*A*_(*σ, σ*′) is the time-varying strength of the constraint. The target distance between neighboring apical domains is constant throughout the simulations and set to 3 pixels, a value much shorter than the initial width of the cells (10 pixels). Initially the constraint *λ*_*A*_(*σ, σ*′), which we interpret here as the combined strength of apical cytoskeletal activity between cell pairs, is set very low (*λ*_*A*_ = 20), so that there is no effect on the tissue or individual cell shapes. A similar form (*H*_*Basal links*_) is used for neighbouring basal domains with a fixed *λ*_*B*_ = 100. This term ensures that the top part of the tissue stays together once the apical compartments have separated.

Above the basal side of cells, there is a collection of domains representing the ectoderm and fibronectin- and laminin-rich extracellular matrix that forms a basal lamina [26]. These domains have a volume constraint and liable adhesion with the basal side of the epithelial cells that helps to maintain the alignment of the cells below them, while allowing some upward/downward movement of the cells. Below their apical sides, cells are adjacent to a loose mesenchyme (mainly corresponding to the future “core” of the somite, or somitocoel) which we choose to model as a single domain with no volume constraint and labile adhesion to the cells.

At the rostral and caudal ends of the row of cells, we include a single immotile cell, which does not change in shape or properties over time. Between them there are 115 to 205 cells, depending on the conditions being tested in the simulations. Apart from the fixed cells that make the AP boundaries, all cells have identical properties and are allowed to change shape and properties during the course of the simulation.

Initially all cells are in an inactive state, without exerting significant forces on each other. We define a ‘wave of constriction activation’ which moves through the tissue from the rostral end at a fixed speed *W* (in units of cell/MCS), so that the next caudal cell is activated 1/*W* MCS after the lastly activated cell. Apical constriction results from a linear increase in *λ*_*A*_ between a pair of activated cells. This variable varies from an initial value of 20 up to 600 at a specific build-up rate *Λ* = *dλ*_*A*_/*dt* (in units of 1/MCS). As *λ*_*A*_ increases, the tension between a pair of cell’s apical domains is:

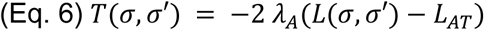

When this tension value exceeds a maximum threshold *T*_*break*_ = −7500 the link between the apical domains of neighboring cells is broken, resulting in the apical separation between the cell pairs.

Variation in cell aspect ratios was implemented by adjusting the internal distance constraints between domains in Eq. 3. We also choose to keep the total sum of all domain target volumes constant and adjust the initial width and length of the cells to satisfy the internal distance constraints.

### Reference Simulation Parameters

Our simulations have 4 parameters, which we will vary systematically in this study:

1. *the wave speed W*, at which the constriction front travels from the rostral to the caudal end of the tissue. The position of the front determines when and where cell pairs start increasing *λ*_*A*_;
2. *the rate of increase of apical contractility* (*build-up rate*) *Λ* = *dλ*/*dt*, which determines how fast the cell pair *λ*_*A*_ increases once activated;
3. *the apical tension breaking threshold T*_*break*_, which determines the tension (Eq. 6) at which apical links between neighboring cells break (Eq. 5); and
4. *the cell aspect ratio AR*, which defines how elongated the cells are at the beginning of the simulation.

The reference values of all 4 parameters can be found in (Table 1). Rest of the simulation parameters are listed in the Supplementary Materials.

### Metrics

To analyze the behavior of our model with respect to variation in key parameters we define and measure the following metrics:

1. *Average Segment Size* < *S* >: defined as the mean number of cells within each cohort in our simulations. We measure the sizes both during the course of the simulation - by taking into account the first occurrence of a separation located caudal to the caudal-most segment - and at the end of the simulation in order to assess the presence of any splitting events.
2. *Average Distance* < *D* >: defined as the distance between the last activated cell pair and the last formed boundary of apical link breakage. This metric is measured during the course of the simulation and is presented in units of the average segment size < *S* > for the corresponding set of parameters.
3. *Average Segmentation Time* < *τ* >: defined as the time (in MCS units) elapsed between the appearance of two consecutive boundaries.

To avoid boundary effects, all parameters are measured after the appearance of the first segmentation and before a distance of 3 < *S* > from the last formed segment from the rostral boundary.

## ACKNOWLEDGMENTS

The authors would like to thank Dr. James P. Sluka and Ellen M Quardokus for helpful discussions. This work was supported by NIH grants R01 GM076992, U01 GM111243 and R01 GM077138.

