## Supplemental Material for "A Mechanical Model of Early Somite Segmentation"

### SUPPLEMENTARY FIGURES AND TABLES

| Parameter | Name | Values |
|---|---|---|
| $T_{break}$ | Threshold breaking tension | 10 |
| $W$ | Frequency of spring activation (wave speed) | 1 – 5 |
| $\Lambda$ | Rate of increase of spring constant (build-up rate) | 1 – 2 |
| $k_0$ | Initial strength of spring constraint | 0 – 20 |
| $k_{max}$ | Maximum spring constant | 5 - ∞ |
| $x_0$ | Resting length of spring | 0 - 0.3 |

| | | |
|---|---|---|
| $d$ | Initial distance between springs | 1 |

**Supplemental Table 1 - List of simulation parameters for the elastic model**

| **Simulation objects** | **Properties and Behaviors** |
|---|---|
| Cell basal compartment | Basal compartments exhibit preferential labile adhesion towards other neighboring basal compartments and E-ECM cells. Neighboring compartments are also connected by spring based constraints. http://purl.obolibrary.org/obo/FMA_74542 |
| Cell core/lateral compartment | Core/Lateral compartments only exhibit labile adhesion and adhere to other neighboring core/lateral compartments with no other preference for any of the other compartments. http://purl.obolibrary.org/obo/FMA_30332 |
| Apical compartment | Apical compartments prefer non-modelled medium domains to adhere to and don’t exhibit any other labile adhesion preferences. Neighboring apical domains are also connected by spring based distance constraints between their center of masses. |

| | |
|---|---|
| | They contract as a response to increased contractility strength of their connection to other apical domains. http://purl.obolibrary.org/obo/FMA_74541 |
| E-ECM cell | E-ECM cells provide a boundary for the simulation and exhibit preferential labile adhesion towards cell basal compartments. http://purl.obolibrary.org/obo/FMA_69070, http://purl.obolibrary.org/obo/FMA_9672 |
| Medium | Medium domain represents all extracellular space and mesenchymal pre-somitic cells that lie ventral to the dorsal epithelium. http://purl.obolibrary.org/obo/FMA_70022 |
| Wall | Wall cells represent an immotile physical boundary on the rostral-caudal sides of the simulation. They don’t exhibit any labile adhesion preferences to the modelled cells but prefer the extracellular space. http://semanticscience.org/resource/SIO_000022.rdf |

**Supplemental Table 2 - Biological components and processes**

A modeled epithelial cell (http://purl.obolibrary.org/obo/FMA_66768) comprises of three compartments - Basal, Core/Lateral and apical. ECM material present near the ectoderm is represented by E-ECM cells. Non-modelled space in the simulation is represented by Medium cells. Immotile Wall cells demarcate simulation boundaries

| Parameter | Name | Value |
| --- | --- | --- |
| $\lambda_V$ | Strength of volume constraint | 10 |
| $V_T$ | Compartment target volume | Basal: 40 pixel$^3$<br>Lateral: 120 pixel$^3$<br>Apical: 40 pixel$^3$<br>E-CM: 70 pixel$^3$ |
| $\lambda_A$ | Strength of apical contractility | Variable, time/spatial dependent range is 20-600 |
| $L_{AT}$ | Target distance between neighboring apical domains | 3 pixels |
| $\lambda_B$ | Strength of basal-basal links | 100 |
| $L_{BT}$ | Target distance between neighboring basal domains | 10 pixels |
| $\lambda_I$ | Strength of internal links | 50 |
| $L_I$ | Target distance between internal domains | Basal-Apical domain : 16 pixels<br>Basal-Lateral domain : 8 pixels<br>Apical-Lateral domain : 8 pixels |

| | | |
|---|---|---|
| $T_B$ | CPM noise parameter | 60 |
| $n_{pixel\ copy}$ | Neighboring distance of pixel copies | 2 |
| $n_{contact}$ | Neighboring distance of contact energies | 4 |

**Supplemental Table 3 - List of simulation parameters**

This supplementary table complements Table 1 in the main text.

| **Adhesion Parameters** | Basal | Apical | Core/Lateral | E-ECM | Medium | Wall |
|---|---|---|---|---|---|---|
| Basal | 83.4 | 100.9 | 100.9 | 80.6 | 100.9 | 100.9 |
| Apical | | 100.9 | 100.9 | 100.9 | 80.6 | 100.9 |
| Core/Lateral | | | 83.4 | 100.9 | 100.9 | 100.9 |
| E-ECM | | | | 83.4 | 100.9 | 100.9 |
| Medium | | | | | 100.9 | 69.4 |
| Wall | | | | | | 0 |

**Supplemental Table 4 - Adhesion energy values between modelled cell compartments**

Note that contact energies in CPM/GGH models are symmetric, so the bottom half of the table is not shown.

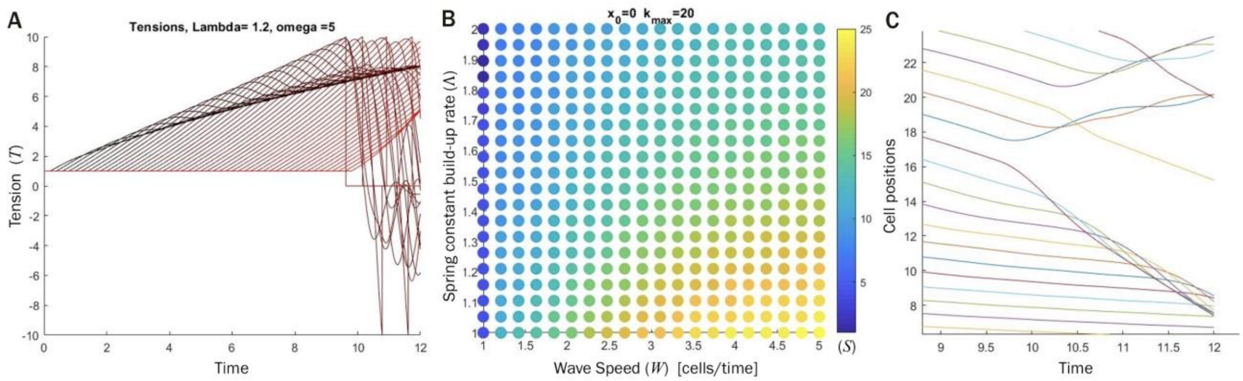


**Supplemental Figure 1 - Linear elastic model results**

(**A**) Time series of cell pair tension values. Each line is the tension of a cell pair (spring) with the black line being the first spring in the chain, and redder lines for each subsequent spring as distance down the chain increases. (**B**) Parameter space diagram for first segment size. Each dot corresponds to set of simulations with a different combination of the parameters $\Lambda$ and $W$ and are colored according to the average segment size (see bar on the right). (**C**) Cell center positions over time at around the moment of the first break (at time ~10), showing subsequent overlap of cell positions.

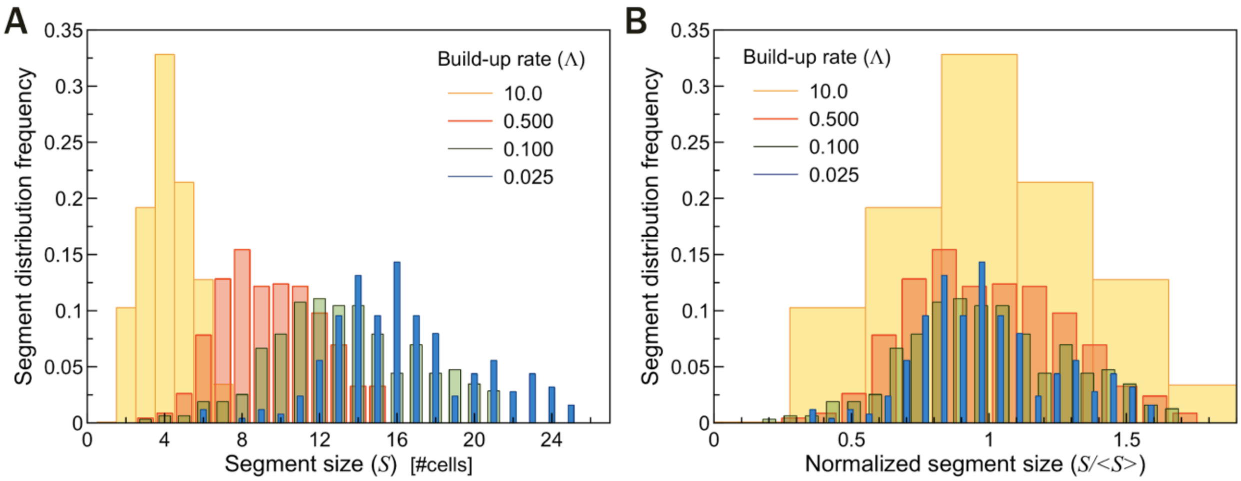

**Supplemental Figure 2 - Segment size distributions for simultaneous increase in contractility and fixed boundary conditions**

(**A**-**B**) Histogram of distribution of segment sizes (**A**) and normalized segment sizes (**B**) for different build-up rates of apical contractility for simulations with fixed cells at each end of the monolayer.

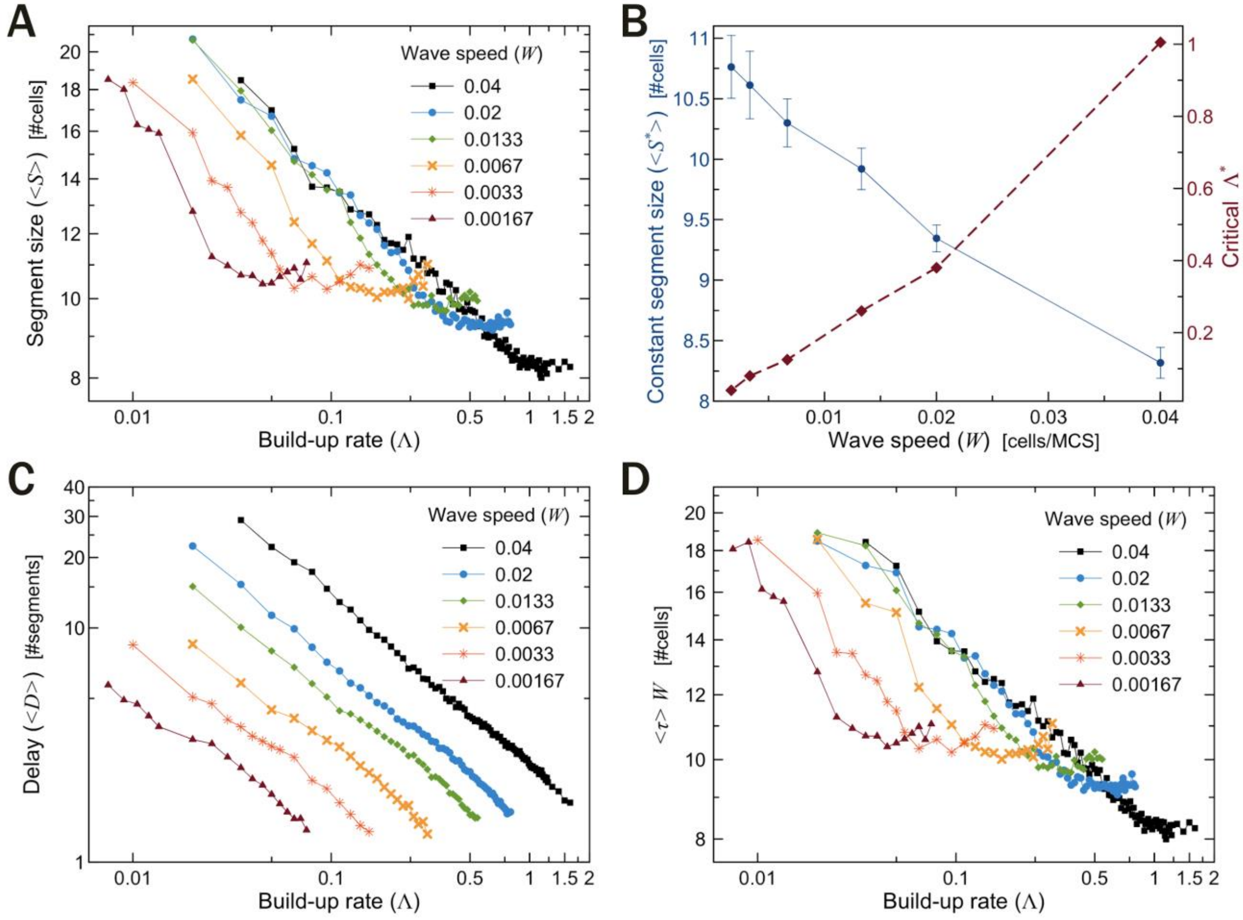


**Supplemental Figure 3 - Segmentation as a function of apical contractility build-up rate**

(**A**) Average segment size $< S >$ as a function of build-up rate of apical contractility $\Lambda$. Segment size decreases logarithmically with build-up rate but stays constant for higher values of $\Lambda$. (**B**) Constant segment sizes $< S^* >$ (blue solid line) and critical build-up rates $\Lambda^*$ (red dashed line) as functions of $W$. (**C**) Distance between activation and boundary formation $< D >$ decreases linearly with $\Lambda$. (**D**) Average segmentation time $< \tau >$ as a function of $\Lambda$. Each individual curve has the same shape as the equivalent

$< S >$ vs. $\Lambda$ curves in (**A**). (**A**,**C**,**D**) Each line shows a qualitative similar behavior for different base values of the wave speed ($W$).

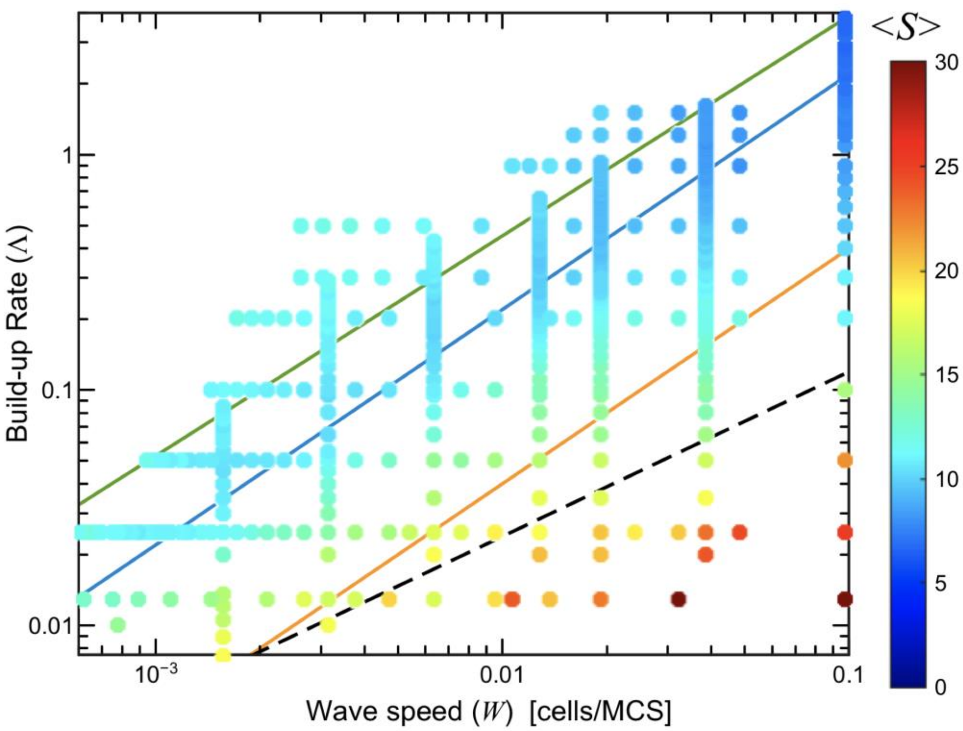


**Supplemental Figure 4 - Parameter space diagram for average segment sizes**

Each dot corresponds to set of simulations with a different combination of the parameters $\Lambda$ and $W$ and are colored according to the average segment size (see bar on the right). Diagonal lines correspond to the boundaries as defined in Figure 5C.

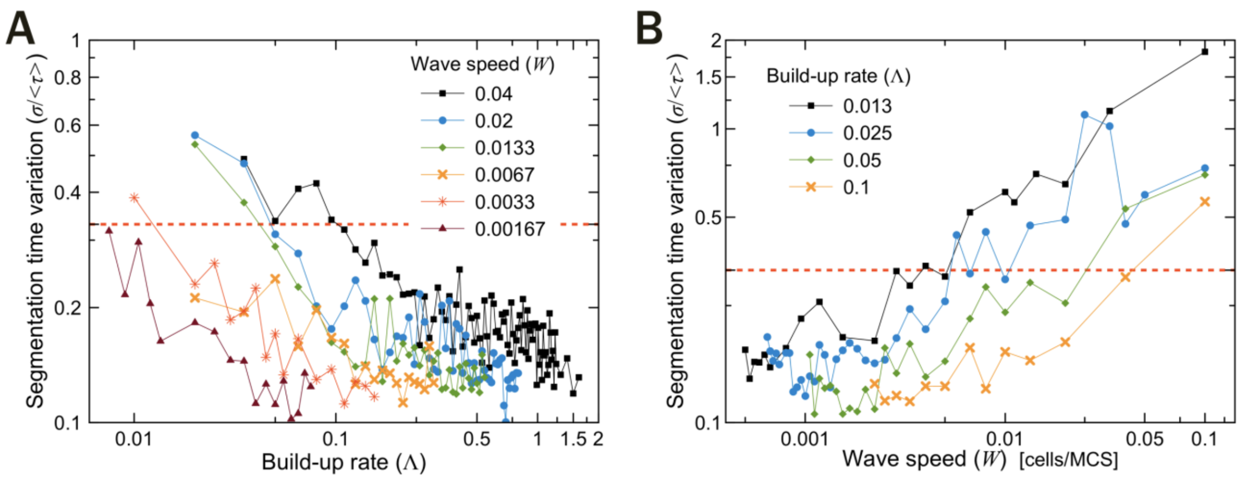


**Supplemental Figure 5 - Segmentation time variation**

(**A**) Segmentation time variation (std/mean) as a function of build-up rate of apical cytoskeletal activity $\Lambda$.

(**B**) Segmentation time variation as a function of wave speed $W$. (**A**,**B**) Dashed red lines at $\sigma/<\tau>= 0.33$ indicate a threshold that gives equivalent criteria to distinguish between regular and irregular segment sizes in Figure 5C.

`
